# Seed dispersal effectiveness and plant-bird interaction networks in the Tehuacán Valley

**DOI:** 10.1101/2024.11.12.623176

**Authors:** Ana María Contreras-González, Mónica Beatriz González-Montes, E. Jair Ortega-Jiménez, Pamela Herrera-Serrano, Francisco Alberto Rivera-Ortiz, Alfonso Valiente-Banuet

**Author notes:** Corresponding Author Email address (AMCG). These authors contributed equally to this work (Mónica Beatriz González-Montes, E. Jair Ortega-Jiménez). These authors also contributed equally to this work.

## Abstract

The maintenance of biodiversity in communities is mediated by interactions that influence community structure, including seed dispersal. The aim of this study was to evaluate the seed dispersal effectiveness of fruit-eating bird species in the Tehuacán Valley and assess the extinction risk of both bird and plant species through interaction networks. From June 2022 to July 2024, we conducted monthly observations of bird foraging on 12 plant species in San Juan Raya and 13 plant species in Zapotitlán Salinas. We calculated the seed dispersal effectiveness and constructed quantitative and qualitative interaction networks for both locations. For both sites, we found that few birds were effective seed dispersers. The most effective dispersers in San Juan Raya were *Melanerpes hypopolius*, *Toxostoma curvirostre*, and *Phainopepla nitens*, while in Zapotitlán Salinas, the main dispersers were *Mimus polyglottos*, *T. curvirostre*, and *P. nitens*. Through qualitative analysis of the interaction networks, we determined that plant communities in both locations are moderately fragile. Quantitative network analyses revealed that plants in Zapotitlán Salinas are particularly vulnerable, as most species could face local extinction if they lose their disperser species, potentially disrupting the community structure. In contrast, bird species exhibited a lower degree of specialization. Thus, even if certain plant species are lost, birds could continue to interact with other species or migrate to areas with a greater availability of resources. The extraction of plant species, many of which are used for firewood, could negatively impact seed dispersal, as these plants serve as facilitators for many of the species studied. This could reduce the number of available sites where birds can deposit seeds, decreasing the likelihood of seed germination and the establishment of new individuals, which, in turn, may compromise community structure.

## Introduction

The maintenance of biodiversity in communities is mediated by positive and negative biotic interactions, which influence community structure [1–3]. Positive interactions, such as mutualism, are crucial for community functioning as they benefit both interacting groups. Many plant species are involved in mutualistic interactions with animals, in which animals provide essential functions for the plant’s reproduction and recruitment, while plants provide energy-rich food resources to various animal species [4–6]. Both seed dispersal (which benefits plants) and antagonistic interactions like seed predation affect the demography of the interacting species, either facilitating or limiting the recruitment of new individuals, and thus play an important role in organismal diversity community [5, 7–9].

During the seed removal phase from plants with animal-dispersed syndromes (zoochory), either dispersal or pre-dispersal predation occurs when fruits or seeds are still on the parent plant. Various organisms, such as ants, birds, bats, rodents, and other mammals, participate in this phase [2, 8, 10, 11]. Seed dispersers benefit plants because they help seeds escape density-dependent mortality caused by pathogens or herbivores beneath the parent plant [2, 8]. Seed dispersal effectiveness (SDE) is an index that assesses the benefit or impact of a seed dispersing species on plant fitness and is composed of both quantity and quality components. Factors such as the number of visits, the number of seeds consumed per visit, the abundance of dispersers, and the locations where seeds are deposited factor into SDE [12, 13]. In arid and semi-arid zones, the availability of suitable sites for seedling establishment also contributes to SDE [14, 15], as many plants require that their seeds be deposited under perennial plants known as nurse plants, where lower solar radiation, transpiration, and temperature, along with higher soil moisture and nutrients, promote seedling establishment [16]. In stressful environments, it is crucial for animal-dispersed seeds to be placed under the canopy of other plants, far from the parent plant, to avoid post-dispersal predation, herbivory, or intraspecific competition. In these zones, bats and birds are the main dispersers, as they are capable of moving seeds over long distances in a short time and placing them in suitable sites, enhancing germination and establishment [2, 8, 17–19].

Birds, by consuming seeds, can either benefit or harm them. In some cases, the scarification of seeds as they pass through the bird’s digestive tract increases germination success, but in other cases, the seeds are destroyed, in which cases the bird acts as a seed predator [4, 8]. Thus, the probability of seeds surviving their passage through the birds’ digestive tract is another key factor in SDE [10].

Most studies of seed removal in arid zones of central Mexico have focused on cacti, with few addressing trees and shrubs. The genera *Melanerpes*, *Myiarchus*, *Mimus*, *Toxostoma*, and *Phainopepla* have been described as effective seed dispersers of cactus species [18, 20–22]. However, it is necessary to expand the evaluation of SDE to include more plant species to achieve better community representation in arid zones of central Mexico.

When used to represent biotic interactions, interaction networks can help to understand community functioning by studying the connectivity between different species. These networks identify key species that maintain community stability [12], as well as revealing the vulnerability of the community to species loss (local extinction), which can trigger cascading effects [23].

In Mexico, the extraction of wild plant specimens for producing artisanal beverages like mezcal and pulque has had a significant impact, as different species are harvested from the wild. Moreover, mezcal production requires considerable firewood sourced from perennial woody plants [24, 25].The loss of species may cause ecosystem collapse by disrupting facilitation, pollination, and seed dispersal networks [26], thus altering community structure and functioning [27].

In this study, we therefore evaluated the impact of frugivorous birds during the pre-dispersal seed removal phase in perennial plant species of the Tehuacán Valley in a community context. We used interaction networks to determine the importance of connectivity between frugivorous birds and plants in the studied communities. We expected that the most abundant birds, which remove the greatest number of seeds and deposit them in suitable sites for germination and establishment, would show the highest SDE values and be the most connected species in the interaction networks, playing a crucial role in the studied communities. We also anticipated that the loss of plant species that provide resources to frugivorous birds, due to the extraction of wild agaves and firewood for mezcal production, would affect the connectivity of plant-frugivore bird networks, impacting the structure of xerophilous shrubland communities in the Tehuacán Valley. Specifically, we aimed to evaluate the effectiveness of fruit-eating bird dispersers in the xerophilous shrubland of the Tehuacán Valley, identify key bird species in seed dispersal, and, through interaction networks, determine the degree of connectivity, nestedness, specialization, and robustness, to assess the extinction risk of species within the community structure.

## Materials and Methods

### Study Area

Fieldwork was conducted at two locations in the Tehuacán Valley within the Tehuacán-Cuicatlán Biosphere Reserve, in the state of Puebla, Mexico: San Juan Raya (SJR) and Zapotitlán Salinas (ZS).

The SJR location is situated between coordinates 18° 18’ 08" - 18° 17’ 13.26" N and 97° 39’ 01.24" - 97° 38’ 26" W at an altitude of 1,840 meters above sea level. It has a semi-desert climate with an average temperature of 21°C and mean annual precipitation of 380 mm. The vegetation is dominated by xerophytic scrub, with *Bouteloua gracilis* and *Acacia subungulata* as the dominant species, along with individuals of the family *Boraginaceae* [28, 29].

The ZS location is situated between coordinates 18° 19’ 51.42" - 18° 19’ 18.37" N and 97° 27’ 36.58" - 97° 26’ 23.32" W at 1,450 meters above sea level. It has a semi-arid climate and summer rains [30] (García, 2004). The annual mean temperature ranges between 18°C and 22°C, with annual precipitation between 380 and 400 mm. The predominant vegetation is xerophytic scrub, with *Mimosa luisana*, *Parkinsonia praecox*, and *Senna wislizenii* as dominant species [18, 29].

### Fieldwork

To collect data on pre-dispersal seed removal for various plant species and provide a community-wide perspective, the study was conducted from June 2022 to August 2024 through monthly sampling sessions, each lasting two days at each site. We calculated the Seed Dispersal Effectiveness (SDE) of birds feeding on 12 plant species with ripe fruits in SJR and 13 plant species with ripe fruits in ZS.

During the monthly samplings, both the quantity and quality components of SDE were assessed [31, 32]. The quantity component for each bird species was estimated by multiplying the number of individuals, the number of seeds consumed, foraging time, visit frequency, relative abundance, and the number of records of each bird species feeding on each plant species. The quality component was calculated by multiplying the probability that birds deposited the seeds under perennial plants by the probability of seed survival after passing through the birds’ digestive tract [31], which was obtained from previous studies (S1 Appendix 1).

#### Interaction Records

To collect data on pre-dispersal seed removal, focal observations were conducted at designated observation points. Between three and seven observers were positioned 10 to 15 meters away from the focal plants to avoid disturbing bird activity. Observations were made on plants with ripe fruits, so the number of focal individuals ranged from one to eight. Using binoculars, observers recorded the following data: bird species feeding on the fruits, part of the fruit consumed (seed and/or pulp), number of fruits/seeds consumed, visit frequency, foraging time, number of birds feeding, and the locations birds moved to after foraging. Observations were carried out during periods of peak bird activity, from sunrise until 9:00 AM and in the afternoon from 5:00 PM until dusk [20, 21, 33, 34]. Bird species were determined using bird field guides [35, 36]. To calculate the number of seeds removed by birds from fruits containing numerous seeds, at least 10 ripe fruits from each plant species were collected, and the average number of seeds per fruit was counted. The number of seeds removed by birds during each visit was then estimated by considering the number of pecks and the average number of seeds per fruit [21].

#### Bird abundance

To determine the relative abundance of frugivorous bird species involved in plant-bird interactions at each study site, the fixed-radius point count technique (25 m) was used. Fourteen counting points were established along two 4-kilometer transects, with a 250 m distance between points. These points were monitored monthly from dawn until 9:30 a.m. Two observers remained at each point for 10 minutes, recording bird species and the number of individuals observed. The counting points were revisited for two consecutive days, reversing the direction of the transects each day [37, 38].

#### Probability of seed placement in suitable sites

During foraging observations, the frequency with which birds visited trees, shrubs, columnar cacti, species of the genera Yucca or Beaucarnea, or the ground after foraging was recorded. This frequency was divided by the total number of visits to all microsites where birds perched after foraging [20].

#### Probability of seed germination

We used previously published data to estimate the percentage of seed germination after passing through the birds digestive tract [31]. Due to the lack of specific information for all plant species studied, average germination rates were calculated for each bird species based on data from similar plants described in the literature. For bird species without available data, the germination percentage reported for birds of the same genus or family was used (S1 Appendix 1).

Finally, the quantity component for each bird species for each study site was calculated as follows: Quantity component = [(relative frequency of visits) x (number of seeds removed per visit) x (number of feeding individuals) x (foraging time) x (relative abundance of each bird species observed feeding during the month) x (number of feeding records for each bird species consuming fruits)] [21, 31]. The quality component for birds was calculated as follows: Quality component = (probability of seed germination after passing through the bird’s digestive tract) x (probability of seed placement in the microsites described above) [31].

### Data analysis

To analyze the data collected from number or records and number of seed removed for each plant species, generalized linear models (GLM) with a Poisson distribution were applied to evaluate whether there were significant differences between plant species at each study site. Also, GLM with a Poisson distribution were used to evaluate whether there were significant differences between the bird species at each study site in data from number of feeding records, the number of seeds removed by different bird species feeding on each observed plant, visit frequency, foraging time, of different bird species at each study site [39].

Additionally, the Chi-square test (*X²*) was used to determine if there were significant differences in the consumption of ripe and unripe fruits, as well as to evaluate differences in the visit frequencies of frugivorous bird species to the plant species observed. The Chi-square test was also used to assess whether there were significant differences in the number of visits from bird species across different microsites after foraging at each study site [40].

To analyze the plant-bird interaction patterns in seed removal, both qualitative and quantitative interaction networks were constructed for each site. For the quantitative networks, SDE values were used. The qualitative interaction networks were based on records of bird species consuming seeds from 13 plant species and 23 bird species observed in SJR. For the ZS site, records from 15 plant species and 23 bird species were used. In both cases, parameters such as nestedness, connectivity, specialization, and robustness were calculated.

Nestedness is a measure that describes how species are connected within the network. This parameter is based on temperature T (degree of disorder in the matrix), where T° = 0 indicates a completely nested matrix, and T° = 100 indicates the absence of nestedness [41, 42].

Connectivity measures the percentage of observed links relative to the total possible links within the network. Connectivity values range from 0 to 1; values close to 0 indicate a poorly connected network, while values near 1 suggest a highly connected network [41]. Specialization refers to the number of species each species in the network interacts with. This variable ranges from 0 (no specialization) to 1 (completely specialized) [43]. Finally, robustness is calculated through simulations of species extinction within the network. Robustness values range from 0 to 1, where R = 1 indicates that the network is highly robust to species loss, while R = 0 suggests that the system is fragile under species loss [44]. For the construction of the plant-frugivorous bird interaction networks, the *Bipartite* package in R software was used [45].

### Results

In SJR, we recorded a total of 22 frugivorous bird species feeding on 12 plant species, and in ZS, we observed 21 frugivorous bird species feeding on 13 plant species (S2 Appendix 2). The plant species in SJR with greatest number of bird species feeding on their fruits were *Bursera galeottiana* (11 species), *Myrtillocactus geometrizans* (10 species), and *Yucca periculosa* (eight species) (Fig 1A). The birds feeding on the largest number of plant species were *Melanerpes hypopolius* and *Peucaea mystacalis* (six and five species, respectively) (Fig 2A). In ZS, the plants attracting the greatest number of bird species were *M. geometrizans* (nine species), *Lemaireocereus hollianus* (eight species), and *Stenocereus stellatus* (seven species) (Fig 1B). The bird species feeding on fruits from the greatest number of plants were *Toxostoma curvirostre* (eight species), *M. hypopolius* (seven species), and *Mimus polyglottos* (six species) (Fig 2B). In SJR, M*. geometrizans* had the highest number of birds feeding records (91 records), followed by B*. galeottiana* (76 records). The birds with the most feeding records were *Melanerpes hypopolius* (64 records) and *Myiarchus cinerascens* (51 records) (Fig 2A; *X²* = 720.62, D.F. = 220, *P* < 0.001). In ZS, *L. hollianus* had the highest number of birds feeding records (71 records), followed by *M. geometrizans* (29 records), *Vallesia glabra* (29 records), and *Stenocereus stellatus* (28 records). The bird species with the most feeding records was *M. hypopolius* (77 records), followed by *M. polyglottos* (31 records) (Fig 2B; *X²* = 797.68, D.F. = 240, *P* < 0.001).

**Fig 1.**
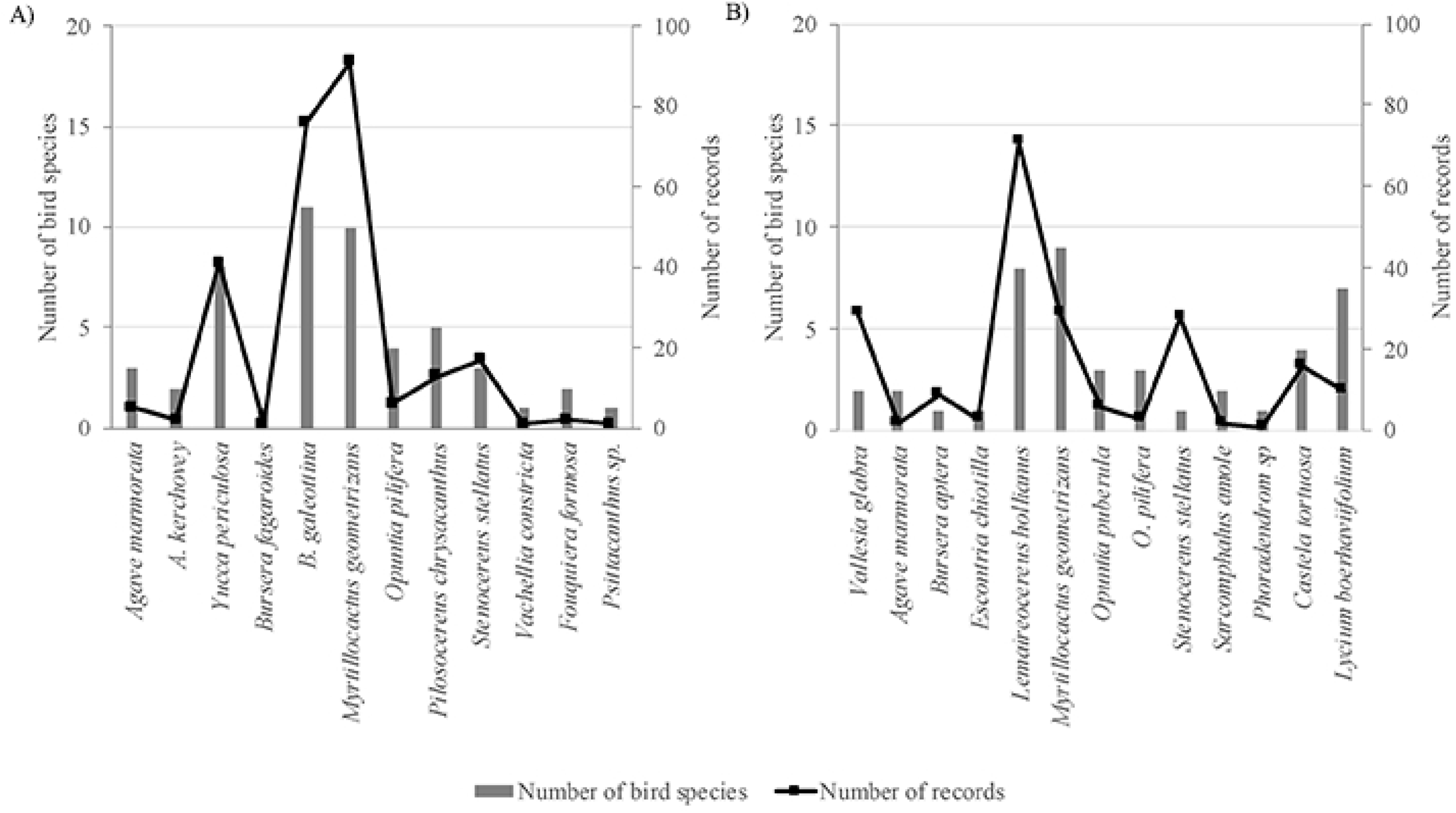
**This is the Fig 1 Title.** This is the Fig 1 Number of bird species recorded foraging on fruits from plant species in San Juan Raya (A) and Zapotitlán Salinas (B).

**Fig 2.**
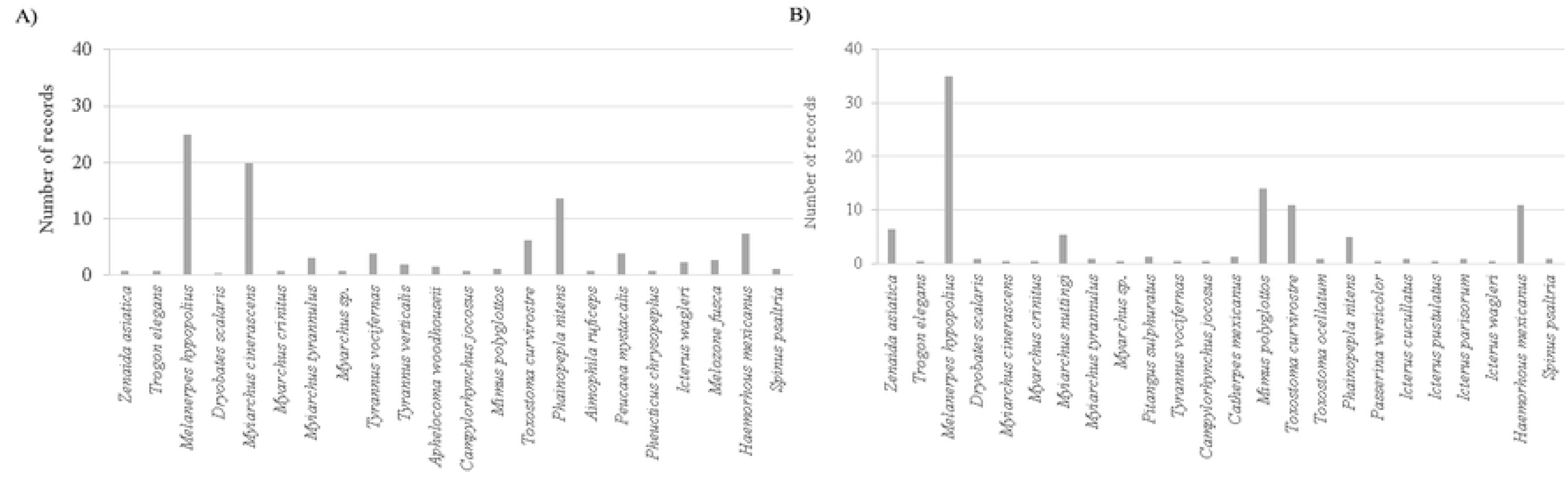
**This is the Fig 2 Title.** This is the Fig 1 Number of records of plant species consumed by birds in San Juan Raya (A) and Zapotitlán Salinas (B).

### Quantity component

Most birds removed seeds from ripe fruits at both sites; however, there were no significant differences in the number of mature and immature seeds consumed by birds (SJR – *X^2^* = 39.8, D.F. = 21, *P* > 0.001; ZS – *X^2^* = 27.60, D.F. = 20, *P* > 0.001). The plants with the highest seed removal in both sites were *M. geometrizans* (SJR – 3,279.64 seeds; ZS – 338 seeds) and *S. stellatus* (SJR – 380 seeds; ZS – 295 seeds). The plants with the lowest seed removal were *Phoradendron sp.* (two seeds), *Psittacanthus sp.* (two seeds), and *Sarcomphalus amole* (three seeds) at both sites (SJR – *X²* = 1,927.8, D.F. =11, *P* < 0.001; ZS – *X²* = 5,502.9, D.F. = 12, *P* < 0.001).

Considering all the studied plants, in SJR, *P. nitens* was the bird species that removed the largest number of seeds (2,387.52 total seeds; 1,193.76 ± 1,185.76 seeds), followed by *Tyrannus vociferans* (595.88 total seeds; 297.94 ± 296.94 seeds) and *M. hypopolius* (510 total seeds; 85 ± 50 seeds). In ZS, *M. hypopolius* removed the largest number of seeds (1,103 total seeds; 180.53 ± 4.87 seeds), followed by *Haemorhous mexicanus* (353 total seeds; 16.80 ± 4.79 seeds) and *T. curvirostre* (238 total seeds; 14 ± 4.07 seeds) (SJR – *X^2^* = 6,434.8, D.F. = 21, *P* < 0.001; ZS – *X^2^* = 1,659.9, D.F. = 20, *P* < 0.001).

In most recorded events, the birds observed feeding on the fruits at both sites generally fed once and then left without returning afterward, because most bird species were observed only once. However, exceptions were observed in SJR, where individuals of *Myiarchus nuttingi*, *M. cinerascens*, and *Melazon fusca* returned multiple times to feed (1.14 ± 0.14, 1.38 ± 0.12, and 1.25 ± 0.12, respectively). Nevertheless, there were no significant differences in the number of visits among the different bird species in this site (*X^2^* = 4.33, D.F. = 21 *P* > 0.001). In ZS, individuals of *M. hypopolius*, *H. mexicanus*, and *T. curvirostre* were recorded returning to feed repeatedly (2.75 ± 0.34, 2.3 ± 0.42, and 1.61 ± 0.4, respectively) (*X^2^* = 53.45, D.F. = 20, *P* < 0.001).

The bird species that spent the most time feeding on the fruits of plants in SJR was *Tyrannus verticalis* (408.75 ± 184.46 seconds), followed by *M. hypopolius* (169.22 ± 54.18 seconds) (*X²* = 5748.6, D.F. = 21, *P* < 0.001). In ZS, the longest recorded feeding time was for *Myiarchus tyrannulus*, although it was observed only once (241 seconds). Additionally, *Myiarchus nuttingi* (219.87 ± 57.89 seconds) and *M. hypopolius* (196.17 ± 23.71 seconds) also spent amount of time feeding on fruits (*X²* = 14,913, D.F. = 20, *P* < 0.001).

The relative abundance of bird species varied over the months during which seed removal was recorded from different plant species. The bird species with the highest relative abundance in SJR were *M. hypopolius* (0.084 ± 0.014) and *H. mexicanus* (0.075 ± 0.01), while in ZS, *Zenaida asiatica* (0.27 ± 0.03), *M. hypopolius* (0.089 ± 0.01), and *M. poliglottos* (0.087 ± 0.009) showed the highest abundance. However, species such as *Myiarchus crinitus* and *T. verticalis* in SJR, as well as *Pitangus sulfuratus*, *Toxostoma ocellatum*, *Passerina versicolor*, *Icterus parisorum*, and *Spinus psaltria* in ZS, were not recorded in any of the count points, resulting in a relative abundance of zero for these species.

### Quality component

The probability of birds depositing seeds in suitable sites for germination, such as trees or shrubs, in SJR was high for *Trogon elegans*, *M. tyrannulus*, *T. vociferans*, *Aphelocoma woodhouseii*, and *Pheucticus chrysopeplus* (with values of 1) (*X^2^* = 239.90, D.F. = 98, P < 0.001). In ZS, the species with the highest probability of depositing seeds in trees or shrubs were *M. tyrannulus*, *P. sulfuratus*, *P. nitens*, and *P. mystacalis* (with values of 1) (Fig 3; *X^2^* = 101.74, D.F. = 64, P < 0.001).

**Fig 3.**
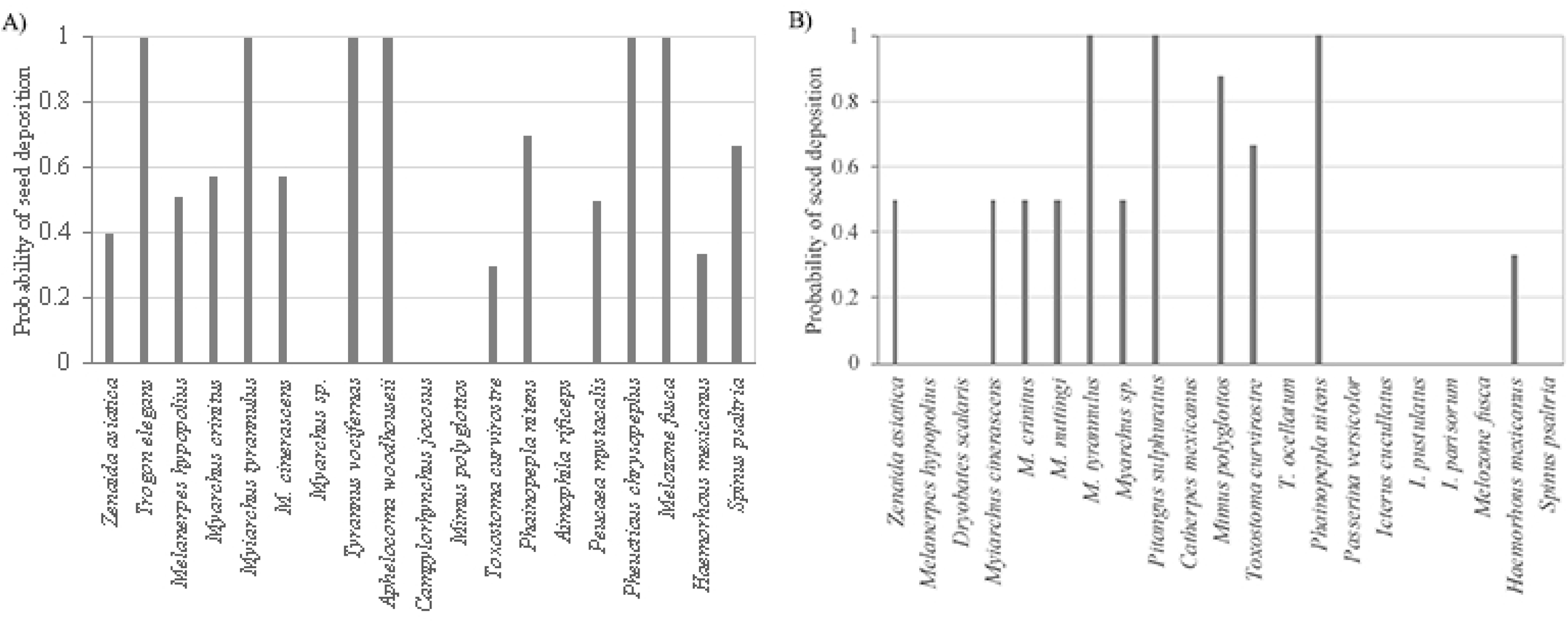
**This is the Fig 3 Title.** This is the Fig 1 Probability of seed deposition by birds in suitable sites for seed germination in San Juan Raya (A) and Zapotitlán Salinas (B).

According to the literature, the species that causes the least damage to seeds during digestion is *T. curvirostre*, with a germination rate of 93 % (± 19.28), followed by *M. polyglottos* (79.33 ± 15.01 % germination), *I. wagleri* (78.12 ± 0.97 % germination), and *M. hypopolius* (64.43 ± 0.97 % germination). Species that completely destroy seeds, resulting in a 0 % germination rate include *Z. asiatica*, *P. mystacalis*, *P. chrysopeplus*, *P. versicolor*, *H. mexicanus*, and *S. psaltria* (S1 Appendix 1).

### Seed Dispersal Effectiveness (SDE)

Birds were found to contribute to seed dispersal of plants to varying degrees. The same bird species could contribute strongly to seed dispersal for some plants and have virtually no contribution for same plant species in different site and others plant species. For example, *M. hypopolius* had a high SDE for *Y. periculosa*, *M. geometrizans*, and *S. stellatus* in SJR, but in ZS, this same plant species had an SDE value of zero. Conversely, *M. polyglottos* showed an SDE value of zero in SJR, yet in ZS, it was one of the birds with the highest SDE values for *M. geometrizans*, *Castela tortuosa*, and *Lycium boerhaviifolium*.

In SJR, *M. hypopolius* had the highest SDE values for *M. geometrizans* (Log 10 SDE = 5.99), *S. stellatus* (Log 10 SDE = 5.22), and *Y. periculosa* (Log 10 SDE = 5.11). Similarly, *P. nitens* also showed high SDE values for *M. geometrizans* (Log 10 SDE = 5.16). In ZS, *T. curvirostre* had the highest SDE values for *L. hollianus* (Log 10 SDE = 5.02) and *S. stellatus* (Log 10 SDE = 4.27), followed by *P. nitens* for *M. geometrizans* (Log 10 SDE = 4.99), *M. polyglottos* for *M. geometrizans* (Log 10 SDE = 4.88) and *C. tortuosa* (Log 10 SDE = 4.35) (Fig 4).

**Fig 4.**
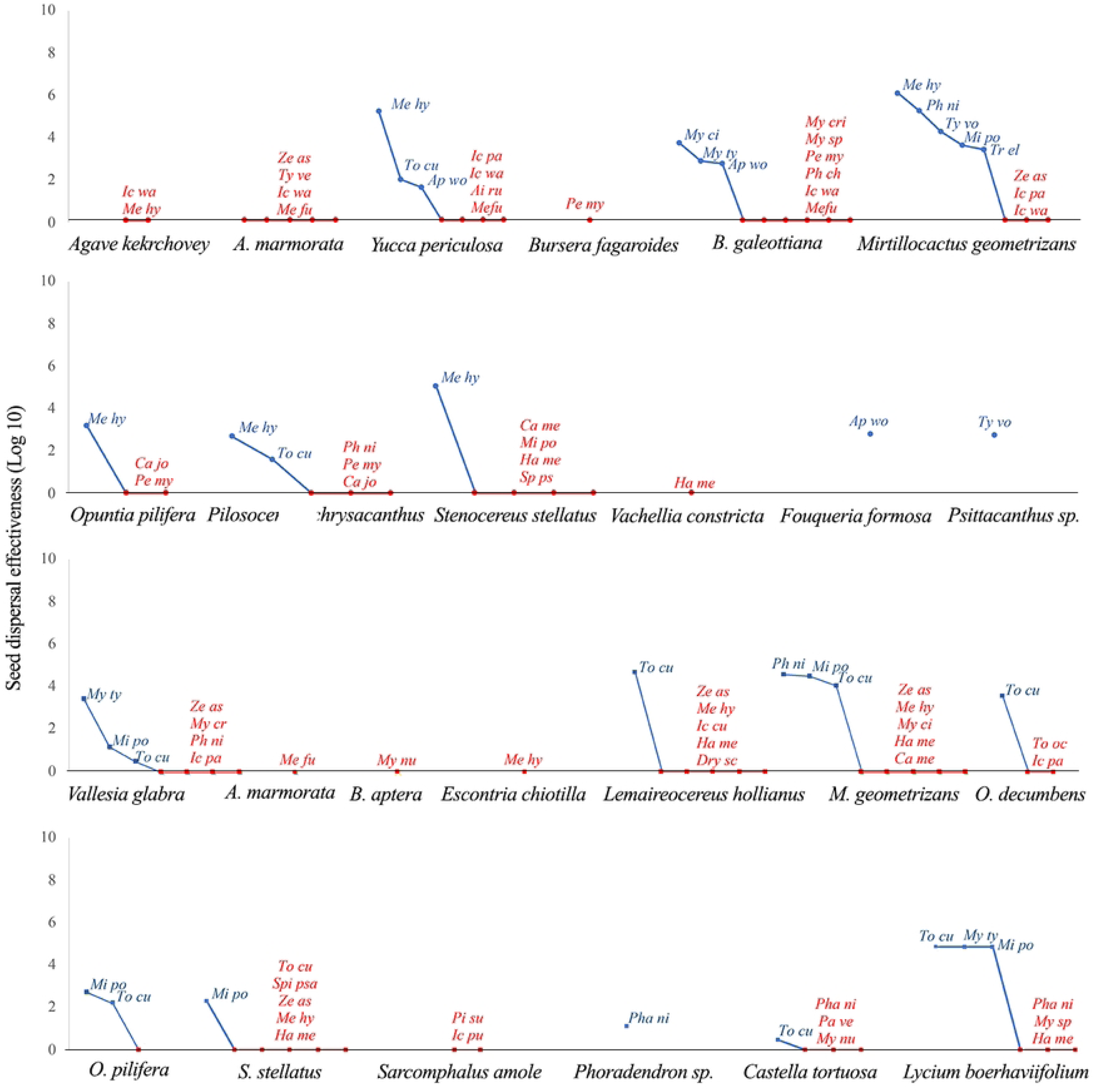
**This is the Fig 4 Title.** This is the Fig 1 Seed dispersal effectiveness by birds consuming fruits in San Juan Raya (circles) and Zapotitlán Salinas (squares). Blue letters represent disperser birds, while red letters represent non-disperser birds. *Ai ru – Aimophila ruficeps, Ap wo - Aphelocoma woodhouseii, Ca jo - Campylorhynchus jocosus, Ca me - Catherpes mexicanus, Drs sc - Dryobates scalaris, Ha me - Haemorhous mexicanus, Ic cu - Icterus cucullatus, Ic pa - Icterus parisorum, Ic pu - Icterus pustulatus, Ic wa - Icterus wagleri, Me hy - Melanerpes hypopolius, Me fu - Melozone fusca, Mi po - Mimus polyglottos, My ci - Myiarchus cinerascens, My crinitus - Myiarchus crinitus, My nu - Myiarchus nuttingi, My sp - Myiarchus sp., My ty - Myiarchus tyrannulus, Pa ve - Passerina versicolor, Pe my - Peucaea mystacalis, Ph ni - Phainopepla nitens, Ph cr - Pheucticus chrysopeplus, Pi su - Pitangus sulphuratus, Sp ps - Spinus psaltria, To cu - Toxostoma curvirostre, To oc - Toxostoma ocellatum, Tr el - Trogon elegans, Ty ve - Tyrannus verticalis, Ty vo - Tyrannus vociferans, Ze as - Zenaida asiatica*

On the other hand, bird species that do not contribute to the seed dispersal of any of the plants they consume include *Z. asiatica*, *Dryobates scalaris*, *M. crinitus*, *M. nuttingi*, *P. sulfuratus*, *T. ocellatum*, *Aimophila ruficeps*, *P. mystacalis*, *P. chrysopeplus*, *P. versicolor*, *I. cuculatus*, *I. pustulatus*, *I. parosorum*, *I. wagleri*, *Melozone fusca*, *H. mexicanus*, and *Spinus psaltria* (Fig 4).

### Network structure

In the qualitative interaction networks, a total of 255 visits from 23 bird species feeding on 13 plant species were recorded in SJR, while 220 visits from 23 bird species foraging on 14 plant species were observed in ZS during the sampling period. The plant species with the highest number of connections in SJR were *B. galeottiana*, *M. geometrizans*, and *Y. periculosa*. The bird species with the most connections were *M. hypopolius*, *P. mystacalis*, and *I. wagleri*. In ZS, the plant species with the most connections were *M. geometrizans*, *L. hollianus*, *V. glabra*, and *L. boerhaviifolium*, while the bird species with the highest number of connections were *T. curvirostre*, *M. polyglottos*, and *M. hypopolius* (Fig 5).

**Fig 5.**
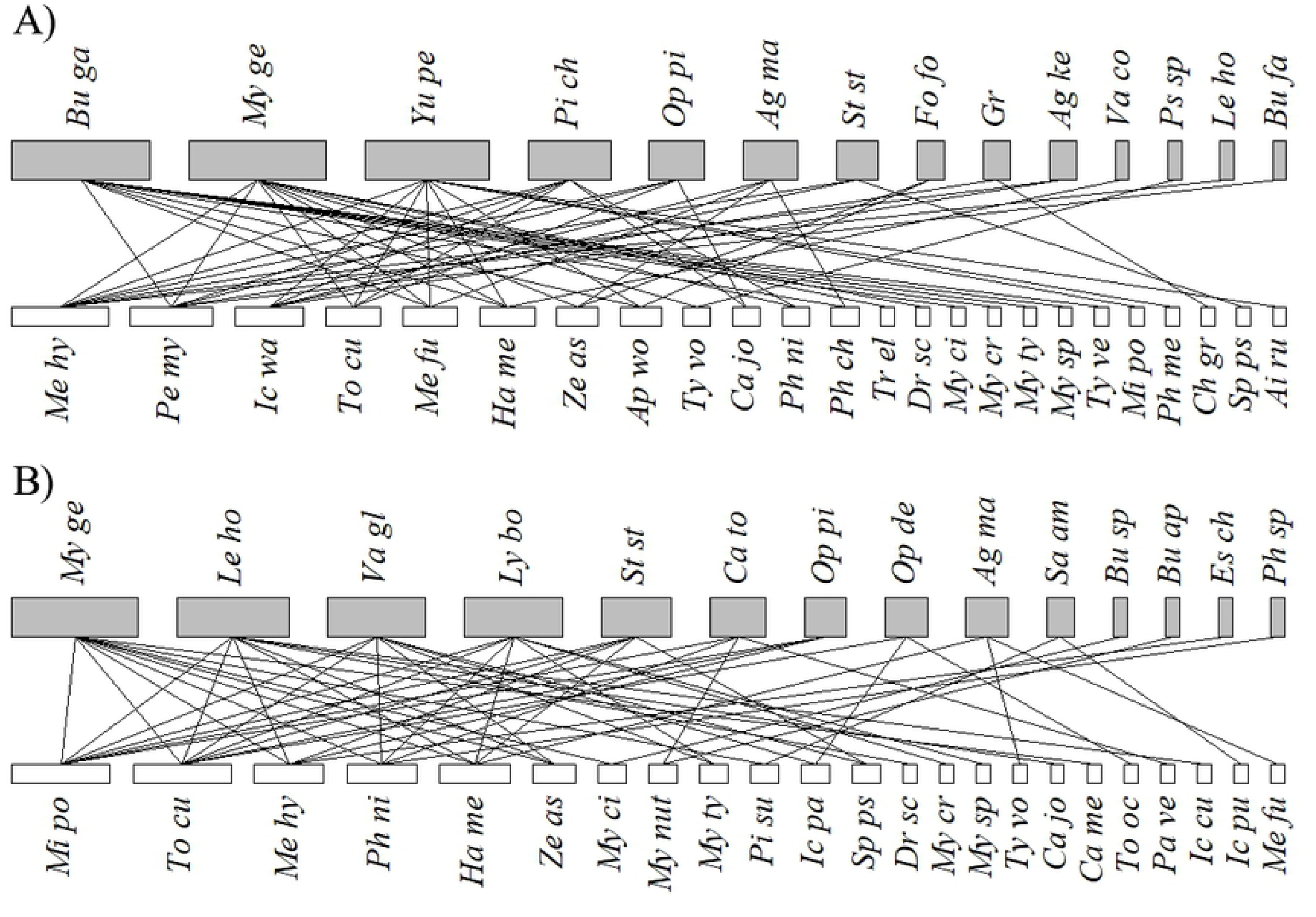
**This is the Fig 5 Title.** This is the Fig 1 Bird-plant visitation networks in the xeric scrubland of the Tehuacán Valley. A) San Juan Raya and B) Zapotitlán Salinas. Gray nodes represent plant species, and white nodes represent bird species. *Ai ru – Aimophila ruficeps, Ap wo - Aphelocoma woodhouseii, Ca jo - Campylorhynchus jocosus, Ca me - Catherpes mexicanus, Ch gr - Chondestes grammacus, Dr sc - Dryobates scalaris, Ha me - Haemorhous mexicanus, Ic cu - Icterus cucullatus, Ic pa - Icterus parisorum, Ic pu - Icterus pustulatus, Ic wa - Icterus wagleri, Me hy - Melanerpes hypopolius, Me fu - Melozone fusca, Mi po - Mimus polyglottos, My ci - Myiarchus cinerascens, My cr - Myiarchus crinitus, My nut - Myiarchus nuttingi, My sp - Myiarchus sp., My ty - Myiarchus tyrannulus, Pa ve - Passerina versicolor, Pe my - Peucaea mystacalis, Ph ni - Phainopepla nitens, Ph ch - Pheucticus chrysopeplus, Ph me - Pheucticus melanocephalus, Pi su - Pitangus sulphuratus, Sp ps - Spinus psaltria, To cu - Toxostoma curvirostre, To oc - Toxostoma ocellatum, Tr el - Trogon elegans, Ty ve - Tyrannus verticalis, Ty vo - Tyrannus vociferans, Ze as - Zenaida asiatica*.

The qualitative plant-bird interaction networks at both sites show low connectance (SJR - 0.17, ZS - 0.17) and high nestedness (SJR - T° = 14.17; ZS - T° = 22.64). Both plant and bird systems at both sites are moderately fragile (plants: SJR-R = 0.58, ZS R = 0.56; birds: SJR-R = 0.68, ZS R = 0.69).

In the quantitative interaction network, low connectance was found in SJR (0.26), while ZS showed medium connectance (0.42). Both sites exhibited high nestedness (SJR - T° = 21; ZS - T° = 21.74). In SJR, plants were moderately vulnerable to extinction risk (R = 0.51), whereas in ZS, the system was more fragile, with most plants losing their interactions, which could lead to extinction (R = 0.38). In contrast, birds showed a lower extinction risk at both sites (SJR R = 0.52, ZS R = 0.68).

## Discussion

Seed dispersal in arid zones plays a crucial role in community structure, as this process ensures that seeds are deposited in suitable locations for successful germination and establishment of new individuals under trees and shrubs. This process influences the spatial arrangement of plants and plant diversity [2, 9].

The role of frugivorous birds in the plant dispersal cycle depends on both quantitative and qualitative components [13, 31, 32, 46]. In this study, seed dispersal effectiveness allowed us to accurately assess birds’ contribution to plant fitness [31, 32]. Additionally, through interaction networks, it was possible to identify whether species loss may affect the structure of the interaction network and, consequently, the community organization [47].

In the xeric scrub of the Tehuacán Valley, a total of 23 species of fruit-eating birds were recorded at each study site. Some of these birds are not exclusively frugivorous but also belong to omnivorous and/or granivorous guilds [48]. Most bird species recorded in this study have previously been described as seed removers for some plant species [18, 21, 22, 49, 50, 51, 52, 53, 54]. However, species like *A. woodhouseii*, *Catherpes mexicanus*, and *T. ocellatum* had not been previously recorded feeding on fruits. The number of bird species feeding on the fruits recorded in this study demonstrates how the frugivorous bird community can concentrate in areas with high resource availability [55].

Additionally, the birds observed in this study do not exclusively feed on fruits with ornithochoric syndrome (red and fleshy fruits) but also take advantage of fruits or seeds with different characteristics, such as white-colored or dry fruits, from which they consume the seeds. This is the case for the seeds of *Agave marmorata*, *A. kerchovey*, *Fouquieria formosa*, and the fruits of *Y. periculosa*, *Vachelia constricta*, *Phoradendron sp.*, and *L. boerhaviifolium*. This is due to seasonal variation in food resources in these environments [56], and fruit choice may depend on the season, phenological availability, crop size, accessibility, fruit size, and shape, as well as the amount of water the birds can obtain from them [57, 58, 59, 60]. Among the species observed at the study sites, other plants produce fruits that contrast with their surroundings and could potentially be dispersed by birds, such as *Polaskia chende*, *Polaskia chichipe*, *Morkillia mexicana*, *Arctostaphylos pungens*, *Peniocereus viperinus*, among others [61, 62]. However, no mature fruits of these species were observed during the sampling.

In the Tehuacán Valley, there are few effective bird dispersers, and these are only effective for certain plant species (*T. elegans*, *M. hypopolius*, *M. cinerascens*, *M. tyrannulus*, *T. vociferans*, *A. woodhouseii*, *T. curvirostre*, and *P. nitens*). In some cases, they do not favor seed dispersal of other plant species. For example, *M. hypopolius* and *M. polygottos* show high SDE values in SJR, while in ZS, SDE values are zero, even though a large number of seeds are removed, and the birds primarily feed on mature fruits. Previous studies also indicate that seeds consumed by these bird species have a high probability of germination [18, 21, 49, 52, 54].

However, it has been described that landscape structure affects seed dispersal patterns [63] (Wang, 2023). In ZS, *M. hypopolius* and *M. polygottos* mainly perch on conspecific individuals or phylogenetically close species after feeding, as densities of up to 1200 individuals/ha of the Cactaceae family have been recorded in this area [18]. This behavior has been previously observed for *M. hypopolius* [18, 20, 21], and the same was observed in this study for *M. polygottos* in ZS. This behavior affects seed dispersal since germination and seedling establishment are low under parental individuals or phylogenetically close species [21, 28, 64], due to the action of pathogens and herbivores that reduce the survival of new individuals [2, 65, 66].

Some bird species, such as *D. scalaris*, *M. cinerascens s*, *M. crinitus*, *M. nuttingi*, *P. sulphuratus*, *T. verticalis*, *T. ocellatum*, *P. nitens*, *I. parisorum*, *I. pustulatus*, and *I. waglerii*, though they do not damage seeds while consuming them, had low or zero relative abundance during the months that they were observed removing seeds, meaning they do not contribute significantly to seed dispersal due to their scarcity at the study sites. Finally, bird species such as *Z. asiatica*, *P. mystacalis*, *P. chrysopeplus*, *P. versicolor*, *M. fusca*, *H. mexicanus*, and *S. psaltria* damage the seeds during their passage through the digestive tract, resulting in non-viable dispersal [18, 21, 22, 52]. Both seed predation and the low abundance of these species contribute to SDE values of zero, negatively impacting plant population dynamics [8, 67]. Although bird predation has sometimes been suggested to regulate plant populations [68], in our study sites, this population dynamic may be affected by the low number of effective bird dispersers, which seems to be related to fruit availability, plant density, and frugivorous bird richness in the community [69, 70, 71]. Furthermore, the extraction of plants used for mezcal production in this part of the reserve [72] (Valiente-Banuet & Verdú, 2007) could have negative effects on seed rain patterns, affecting plant spatial distribution, demography, and landscape structure [32, 73, 74, 75].

Interaction networks in SJR and ZS highlight the organization of the plant-frugivore community, showing that certain bird species connect with various plant species, exhibiting a higher number of links. Among the species with the highest SDE values are *T. curvirostre*, *M. polygottos*, *M. hypopolius*, and *P. nitens*, although seed-predating species such as *P. mystacalis* were also found. We found that generalist bird species tend to interact with generalist plant species; however, there were few cases of specialist interactions, a phenomenon described as rare but with the potential to impact the community [76].

Qualitative and quantitative interaction networks from both sites showed low connectivity, which is common in mutualistic interaction networks, where many bird species are observed feeding on plant fruits [12]. Moreover, the networks showed high nestedness, indicating that specialist species interact with generalist species, creating asymmetry in interaction specificity [47, 77, 78]. Using qualitative networks, we found that both plant systems are moderately fragile, while quantitative networks revealed that the SJR plant system is moderately fragile, but the ZS system is more fragile. In this site, most plants could face extinction if they lose their interactions with birds, with rare and specialist species being the most vulnerable [26, 79, 80]. This would affect plant species richness and density, potentially triggering long-term changes in the plant community structure [81]. Regarding birds, both sites exhibit a lower degree of specialization.

Therefore, if plant species were to be lost, birds could maintain interactions with other plants or migrate to sites with greater availability of food resources, impacting bird species richness [26, 55, 56, 81]. However, the role that these bird species play in the study sites cannot likely be compensated for by other species [73] due to the low redundancy of seed-dispersing birds found in this work, making the studied communities more susceptible [82,83].

In addition to the low number of seed-dispersing birds, the loss of plant species caused by the extraction of species used as firewood for the production of drinks such as mezcal could impact seed dispersal, as many of these plant species facilitate the growth of other plant species [72]. If these facilitative species were to be lost in stressful environments, the plant-plant interactions between facilitative species and those facilitated, which are dispersed by birds (as observed in this study), would also be lost, disrupting the seed dispersal phase in the plant life cycle. This would result in a decrease in suitable sites for germination and establishment of new individuals, altering plant recruitment and potentially leading to community-level consequences, such as local species loss and the loss of ecosystem services, thereby affecting community structure [26, 79, 84].

## Conclusions

The data obtained in this study reveal patterns of interaction between frugivorous birds and plants that provide valuable insights, where in both communities there were few effective dispersers. In most cases, it was observed that generalist species interact with specialist species, although associations between specialist species were also found. The SDE analysis and interaction networks allowed us to identify key species within the communities. The qualitative networks indicated moderate fragility of both communities, while the quantitative network showed that SJR is moderately fragile, while ZS is more fragile, primarily due to the loss of seed dispersal and nurse interactions. Therefore, the extraction of species used could significantly impact these communities. This underlines the importance of approaching species conservation programs from a community perspective.

## Acknowledgements

The authors would like to thank the community authorities of San Juan Raya and Zapotitlán Salinas, Puebla, for their support during the field data collection. We also extend our gratitude to Zeltzin Juárez Vicuña, Jesús Ortega Esquinca, Joshua Abraham Jiménez Reséndiz, Ramón García Romero, Joselyn Giovana Pedraza Ramírez, Ariel Emmanuel Montes de Oca Soria, Katia Vianey Montaño Soto, Arturo Alva Saltillán, Apolo López González, and Juan Miguel Gallegos Rojas for their assistance in the field data collection.

## Supporting information

**S1 Appendix. This is the S1 Appendix Title**. This is the S1 Appendix legend. Percentage of seed germination after passage through the gut tract of birds.

**S2 Appendix 2 This is the S2 Appendix Title.** This is the S2 Appendix legend. Bird species foraging fruits from plant species of San Juan Raya (SJR) and Zapotitlán Salinas (ZS).

**S1 Fig. This is the S1 Fig Title**. This is the S1 Fig legend. Number of bird species recorded foraging on fruits from plant species in San Juan Raya (A) and Zapotitlán Salinas (B).

**S2 Fig. This is the S2 Fig Title**. This is the S2 Fig legend. Number of records of plant species consumed by birds in San Juan Raya (A) and Zapotitlán Salinas (B).

**S3 Fig. This is the S3 Fig Title**. This is the S3 Fig legend. Probability of seed deposition by birds in suitable sites for seed germination in San Juan Raya (A) and Zapotitlán Salinas (B).

**S4 Fig. This is the S4 Fig Title.** This is the S4 Fig legend. Seed dispersal effectiveness by birds consuming fruits in San Juan Raya (circles) and Zapotitlán Salinas (squares). Blue letters represent disperser birds, while red letters represent non-disperser birds. *Ai ru – Aimophila ruficeps, Ap wo - Aphelocoma woodhouseii, Ca jo - Campylorhynchus jocosus, Ca me - Catherpes mexicanus, Drs sc - Dryobates scalaris, Ha me - Haemorhous mexicanus, Ic cu - Icterus cucullatus, Ic pa - Icterus parisorum, Ic pu - Icterus pustulatus, Ic wa - Icterus wagleri, Me hy - Melanerpes hypopolius, Me fu - Melozone fusca, Mi po - Mimus polyglottos, My ci - Myiarchus cinerascens, My crinitus - Myiarchus crinitus, My nu - Myiarchus nuttingi, My sp - Myiarchus sp., My ty - Myiarchus tyrannulus, Pa ve - Passerina versicolor, Pe my – Peucaea mystacalis, Ph ni - Phainopepla nitens, Ph cr - Pheucticus chrysopeplus, Pi su - Pitangus sulphuratus, Sp ps - Spinus psaltria, To cu - Toxostoma curvirostre, To oc - Toxostoma ocellatum, Tr el - Trogon elegans, Ty ve - Tyrannus verticalis, Ty vo - Tyrannus vociferans, Ze as - Zenaida asiatica*

**S5 Fig. This is the S5 Fig Title**. This is the S5 Fig legend. Bird-plant visitation networks in the xeric scrubland of the Tehuacán Valley. A) San Juan Raya and B) Zapotitlán Salinas. Gray nodes represent plant species, and white nodes represent bird species. *Ai ru – Aimophila ruficeps, Ap wo - Aphelocoma woodhouseii, Ca jo - Campylorhynchus jocosus, Ca me - Catherpes mexicanus, Ch gr - Chondestes grammacus, Dr sc - Dryobates scalaris, Ha me - Haemorhous mexicanus, Ic cu - Icterus cucullatus, Ic pa - Icterus parisorum, Ic pu - Icterus pustulatus, Ic wa - Icterus wagleri, Me hy - Melanerpes hypopolius, Me fu - Melozone fusca, Mi po - Mimus polyglottos, My ci - Myiarchus cinerascens, My cr - Myiarchus crinitus, My nut - Myiarchus nuttingi, My sp - Myiarchus sp., My ty - Myiarchus tyrannulus, Pa ve - Passerina versicolor, Pe my - Peucaea mystacalis, Ph ni - Phainopepla nitens, Ph ch - Pheucticus chrysopeplus, Ph me - Pheucticus melanocephalus, Pi su - Pitangus sulphuratus, Sp ps - Spinus psaltria, To cu - Toxostoma curvirostre, To oc - Toxostoma ocellatum, Tr el - Trogon elegans, Ty ve - Tyrannus verticalis, Ty vo - Tyrannus vociferans, Ze as - Zenaida asiatica*.

## References

1. Begon M, Townsend C, N., Harper, JL. Communities and Ecosystems. In Ecology from individuals to ecosystems 4th ed. Blackwell Publishing; 2006. pp.550–577. 10.1051/limn/1999024

2. Howe HF, Smallwood J. Ecology of seed dispersal. Ann Rev Ecol Syst. 1982; 13: 201– 228.

3. Kothamasi D, Kiers, ET, van der Heijden, GAM. Community ecology. Processes, models and aplications. Herman V, Peter M editors. Oxford Academic; 2010. 10.1017/cbo9780511574641.004

4. Heleno, RH, Ross G, Everard A, Memmott J, Ramos JA. The role of avian “seed predators” as seed dispersers. Ibis, 2011; 153: 199–203. 10.1111/j.1474-919X.2010.01088.x

5. Howe HF. Seed dispersal by fruit-eating birds and mammals. In Murray DR editor. Seed dispersal North Ryde, Australia. Academic Press; 1986. pp. 417–426.

6. Jordano P. Fruits and frugivory. In Fenner M editor. Seeds: The ecology of regeneration in plant communities 2th. Edition.CABI Publishing; 2000. pp. 125–165

7. Fuentes M. Frugivory, seed dispersal and plant community ecology. Trends Ecol Evol, 2000; 15: 487–488. 10.1016/S0169-5347(00)02031-0

8. Janzen, DH. Seed predation by animals. Ann Rev Ecol Syst. 1971;2: 465–492. DOI:10.1146/annurev.es.02.110171.002341

9. Murray KG, Garcia-CM. Contributions of seed dispersal and demography to recruitment limitation in a Costa Rican cloud forest. In Levey DJ, Silva SW, Galetti M editors. Seed dispersal and frugivory: ecology, evolution and conservation. CAB International; 2002. pp. 323–379.

10. Hulme PE, Benkman CW. Granivory. In Herrera CM, Pellmyr O editors. Plant-animal interactions: evolutionary ecology in tropical and temperate regions Oxford. 2000.

11. van der Pijl. Principles of seed dispersal in higher plants. Springer-Verlag; 1982

12. Jordano P, Vázquez D. Bascompte J. Redes complejas de interacciones mutualistas planta-animal. In Mendel R, Aizen A, Zamora R editors. Ecología y evolución de interacciones planta-animal. Editorial Universitaria; 2009. pp. 17–41.

13. Jordano P, Schupp E. Seed disperser effectiveness: The quantity component and patterns of aeed rain for *Prunus mahaleb*. Ecol Monogr. 2010; 58: 251–269. http://www.jstor.org/stable/2657187

14. Duncan RP, Diez JM, Sullivan JJ, Wangen S, Miller AL. Safe sites, seed supply, and the recruitment function in plant populations. Ecology. 2009; 90: 2129–2138. 10.1890/08-1436.1

15. Harper JL, Williams JT, Sagar GR. The behaviour of seeds in soil: I. The heterogeneity of soil surfaces and its role in determining the establishment of plants from seed. J. Ecol. 1965; 53: 273. 10.2307/2257975

16. Callaway RM. Positive interactions and interdependence in plant communities. Springer; 2007.

17. Connell JH. On the role of natural enemies in preventing competitive exclusion in some marine animals and in rain forest trees. In den Boer PJ, Gradwell G editors. Dynamics of numbers in populations. Wageningen, The Netherlands: Center for Agricultural Publication and Documentation; 1971. pp. 298–312.

18. Godínez-Alvarez, H, Valiente-Banuet A, Rojas-Martínez A. The role of seed dispersers in the population dynamics of the columnar cactus *Neobuxbaumia tetetzo*. Ecology, 2002; 83:(9), 2617–2629. 10.1890/0012-9658(2002)083[2617:TROSDI]2.0.CO;2

19. Massey FP, Press MC, Hartley SE. Have the impacts of insect herbivores on the growth of tropical tree seedlings been underestimated? In Burslem DFR, Pinard MA, Hartley SE editors. Biotic interactions in the tropics: their role in the maintenance of species diversity. Cambridge University Press; 2005. pp. 347–36

20. Castillo LJP. Dispersión biótica de semillas de la cactacea columnar *Neobuxbaumia mezcalaensis* (Bravo) Backeberg en el Valle de Tehuacán, Puebla. PhD Thesis, Universidad Nacional Autónoma de México. 2011.

21. Contreras-González AM, Arizmendi M del C. Pre-dispersal seed predation of the columnar cactus (*Neobuxbaumia tetetzo*, Cactaceae) by birds in central Mexico. Ornitol Neotrop. 2014; 25: 373–387. https://digitalcommons.usf.edu/ornitologia_neotropical/vol25/iss4/1/

22. Ramos-Ordoñez MF, Arizmendi MC. Parthenocarpy, attractiveness and seed predation by birds in *Bursera morelensis*. J Arid Environ. 2011; 75; 757–762. 10.1016/j.jaridenv.2011.04.013

23. Olesen JM, Bascompte J, Dupont YL, Jordano P. The smallest of all worlds: Pollination networks. J Theor Biol. 2006; 240: 270–276. 10.1016/j.jtbi.2005.09.014

24. Pérez DS. Residuos de agave en el proceso de producción de mezcal artesanal en el Valle de Tehuacán-Cuicatlán. Universidad Autónoma del Estado de Morelos, Morelos, México. 202.

25. Torres I, Casas A, Vega E, Martínez-Ramos M, Delgado-Lemus A. Population dynamics and sustainable management of mescal agaves in central Mexico: *Agave potatorum* in the Tehuacán-Cuicatlán Valley. Econ Bot. 2015; 69: 26–41. 10.1007/s12231-014-9295-2

26. Valiente-Banuet A, Verdú M. Human impacts on multiple ecological networks act synergistically to drive ecosystem collapse. Front Ecol Environ. 2013;11: 408–413. 10.1890/130002

27. Wardle DA, Bardgett, RD, Callaway RM, Van der Putten WH. Terrestrial ecosystem responses to species gains and losses. Science. 2011; 332: 1273–1277. https://www.science.org

28. Castillo LJP, Valiente-Banuet A. Species-specificity of nurse plants for the establishment, survivorship, and growth of a columnar cactus. Am J Bot. 2010; 97; 1289–1295. 10.3732/ajb.1000088

29. Ortega-Jiménez EJ. Descripción de la comunidad de aves y su relación con la vegetación del matorral xerófilo en San Juan Raya y el Jardín Botánico Helia Bravo, Puebla. Thesis. Universidad Nacional Autónoma de México. 2024

30. García E. Modificaciones al Sistema Climático Köppen. Instituto de Geografía. 2004 p. 98 p. Retrieved from http://www.igeograf.unam.mx/sigg/utilidades/docs/pdfs/publicaciones/geo_siglo21/serie_lib/modific_al_sis.pdf

31. Schupp EW. Quantity, quality and the effectiveness of seed dispersal by animals. Vegetatio. 1993; 107: 15–29. 10.1007/978-94-011-1749-4_2

32. Schupp EW, Jordano P, Gómez JM. Seed dispersal effectiveness revisited: A conceptual review. New Phytol. 2010; 188: 333–353. 10.1111/j.1469-8137.2010.03402.x

33. Altmann J. Oservational study of behaviour: sampling methods. Behaviour. 1973; 49: 227–266. DOI: 10.1163/156853974x00534

34. Pizo MA. Frugivory and habitat use by fruit-eatin birdsin a fragmented landscape of Southeast Bazil. Ornitol Neotrop. 2004; 15: 117–126. 10.1590/0102-33062016abb0192

35. Howell SNG, Webb S. A guide to the birds of Mexico and North Central America. Oxford University Press. 1995.

36. Kaufman K. Field Guides To Birds of North America. 2005. p. 383.

37. Hutto RL, Pletschet SM. A fixed-radius point count method for nonbreeding and breeding season use. Auk. 1986; 103: 593–602. DOI:10.1093/auk/103.3.593

38. Ralph CJ, Geupel GR, Pyle P, Martin TE, Desante DF, Milá B. Manual de métodos de campo para el monitoreo de aves terrestres. General Technical Report PSW-GTR- 159. Department of Agriculture: 1996. 46 p. http://www.srs.fs.usda.gov/pubs/31462

39. Crawley MJ. The R book. 2007. 10.1080/01621459.1989.10478754

40. Zar JH. Bioestadistical. 2010. 10.1017/CBO9781107415324.004

41. Dormann CF, Gruber B, Fründ J. Introducing the bipartite Package: Analysing Ecological Networks (Vol. 8). 2008. Retrieved from http://erzuli.ss.uci.edu/R.stuff.

42. Lara-Rodríguez NZ, Díaz-Valenzuela R, Martínez-García, V, Mauricio-Lopéz E, Anaid-Díaz S, Valle OI, et al. Redes de interacción colibrí-planta del centro-este de México. Rev Mex Biodivers. 2012; 83; 569–577. 10.22201/ib.20078706e.2012.2.965

43. Blüthgen N, Menzel F, Blüthgen. Measuring specialization in species interaction networks. BMC Ecol, 6: 9 10.1186/1472-6785-6-9

44. Burgos E, Ceva H, Perazzo RPJ, Devoto M, Medan D, Zimmermann M, et al. Why nestedness in mutualistic networks? J Theor Biol. 2007; 249: 307–313. 10.1016/j.jtbi.2007.07.030

45. R Development Core Team. R: a language and environment for statistical computing. Vienna: R Foundation for Statistical Computing. 2024. http://www.R-project.org

46. Calviño-Cancela M, Martín-Herrero J. Effectiveness of a varied assemblage of seed dispersers of a fleshy-fruited plant. Ecology. 2009; 90; 3503–3515. 10.1890/08-1629.1

47. Bascompte J, Jordano P. Plant-animal mutualistic networks: The architecture of biodiversity. Ann Rev Ecol Syst. 2007; 38: 567–593. 10.1146/annurev.ecolsys.38.091206.095818

48. Arizmendi M del C, Espinosa MA. Avifauna de los bosques de cactáceas columnares del Valle de Tehuacán, Puebla. Acta Zool Mex. 1996; 67; 25–46. DOI:10.21829/azm.1996.67671755

49. Almazán-Núñez RC, Alvarez-Alvarez EA, Sierra-Morales P, Rodríguez-Godínez R. Fruit size and structure of zoochorous trees: Identifying drivers for the foraging preferences of fruit-eating birds in a Mexican successional dry forest. Animals. 2021; 11; 10.3390/ani11123343

50. Almazán-Núñez RC, Eguiarte LE, Arizmendi M del C, Corcuera P. *Myiarchus flycatchers* are the primary seed dispersers of *Bursera longipes* in a Mexican dry forest. PeerJ, 2016; 15. 10.7717/peerj.2126

51. Álvarez-Espino R, Ríos-Casanova L, Godínez-Álvarez H. Seed removal in a tropical North American desert: an evaluation of pre- and post-dispersal seed removal in *Stenocereus stellatus*. Plant Biol. 2017; 19: 469–474. 10.1111/plb.12541

52. Pérez-Villafaña M, Valiente-Banuet A. Effectiveness of dispersal of an ornithocorous cactus *Myrtillocactus geometrizans* (Cactaceae) in a patchy environment. Open Biol J. 2009; 2: 101–113. 10.2174/1874196700902010101

53. Dunning JB. Foraging choice in three species of Pipilo (Aves: Passeriformes): a test of the threshold concept. PhD Thesis. The University of Arizona. 1986.

54. García VO. Dispersión biótica de semillas de la cactácea columnar *Stenocereus prinosus* (Otto) F.Buxb. en el Valle de Tehuacán-Puebla, México. Thesis Universidad Nacional Autónoma de México. 2000.

55. Tellería JL, Ramirez A, Pérez-Tris J. Fruit tracking between sites and years by birds in Mediterranean wintering grounds. Ecography. 2008; 31; 381–388. 10.1111/j.0906-7590.2008.05283.x

56. Pavón NP, Briones O. Phenological patterns of nine perennial plants in an intertropical semi-arid Mexican scrub. J Arid Environ. 2001; 49: 265–277. 10.1006/jare.2000.0786

57. Burns KC, Dalen JL. Foliage color contrasts and adaptive fruit color variation in a bird-dispersed plant community. Oikos. 2002; 96: 463–469. 10.1034/j.1600-0706.2002.960308.x

58. Cade TJ, Greenwald LI. Drinking behavior of Mousebirds in the Namib desert, Southern Africa. Auk, 1966; 83: 126–128. 10.2307/4082984

59. Flörchinger M, Braun J, Böhning-Gaese K, Schaefer HM. Fruit size, crop mass, and plant height explain differential fruit choice of primates and birds. Oecologia. 2010; 164: 151– 161. 10.1007/s00442-010-1655-8

60. Smith AD, McWilliams SR. Fruit removal rate depends on neighborhood fruit density, frugivore abundance, and spatial context. Oecologia 2014; 174: 931–942. 10.1007/s00442-013-2834-1

61. Arias TAA, Valverde MT, Reyes SJ. Las plantas de la región de Zapotitlán Salinas, Puebla. Instituto Nacional de Ecología.

62. Téllez VO, Reyes CM, Dávila AP, Gutiérrez GK, Téllez PO, Álvarez ER, et al. Guía ecoturística de las plantas del Valle de Tehuacán-Cuicatlán. In Volks Wagen, UNAM, Millenium Seed Bank Project KEW. 2008.

63. Wang Z, Chen Q, Gu Z, Tang N, Li, N. Effects of landscape features on the structure and function of bird seed dispersal networks in fragmented forests. For Ecol Manag. 2023; 545. 10.1016/j.foreco.2023.121251

64. 10.1016/j.foreco.2023.121251

65. Valiente-Baunet A, Ezcurra E. Shade as a cause of the association between the cactus *Neobuxbaumia tetetzo* and the nurse-plant *Mimosa luisana* in the Tehuacan Valley, Mexico. J Ecol. 1991; 79: 961–971. https://www.jstor.org/stable/2261091

66. Janzen DH. Herbivores and the number of tree species in tropical forests. Am Nat. 1970; 104: 501–528. https://www.jstor.org/stable/2459010

67. Wright SJ. Plant diversity in tropical forests: A review of mechanisms of species coexistence. Oecologia. 2002; 130: 1–14. 10.1007/s004420100809

68. Schupp EW, Fuentes M. Spatial patterns of seed dispersal and the unification of plant population ecology. Ecoscience. 1995; 11: 78. https://www.jstor.org/stable/42900841

69. Renton K. Lilac-crowned Parrot diet and food resource availability: resource tracking by a parrot seed predator. Condor, 2001; 103: 62–69. 10.1093/condor/103.1.62

70. Pizo MA. Seed dispersal and predation in two populations of *Cabralea canjerana* (Meliaceae) in the Atlantic forest of southeastern Brazil. J Trop Ecol. 1997; 13: 559–577. https://www.jstor.org/stable/2560180

71. Snow DW. Evolutionary aspects of fruit-eating by birds. Ibis. 1971; 113: 194–202. 10.1111/j.1474-919X.1971.tb05144.x

72. Wheelwright NT. Fruit-size, gape width, and the diets of fruit-eating birds. Ecology. 1985; 66: 808–818. 10.2307/1940542

73. Valiente-Banuet A, Verdú M. Facilitation can increase the phylogenetic diversity of plant communities. Ecol Lett. 2007; 10: 1029–1036. 10.1111/j.1461-0248.2007.01100.x

74. Loiselle BA, Blake JG. Potential consequences of extinction of frugivorous birds for shrubs of a tropical wet forest. In Levey DJ, Silva WR editors. Seed dispersal and frugivory: ecology, evolution and conservation. CAB International; 2002. pp. 397–406.

75. McConkey KR, Prasad S, Corlett RT, Campos-Arceiz A, Brodie JF, Rogers H, et al. Seed dispersal in changing landscapes. Biol Conserv. 2012; 146: 1–13. 10.1016/j.biocon.2011.09.018

76. Pérez-Méndez N, Jordano P, Valido A. Downsized mutualisms: Consequences of seed dispersers’ body-size reduction for early plant recruitment. Perspec Plant Ecol Evol Syst. 2014; 17: 151–159. DOI:10.1016/j.ppees.2014.12.001

77. Bascompte J, Jordano P, Melián CJ, Olesen JM. The nested assembly of plant-animal mutualistic networks. Proc Nat Acad Sci. 2003; 100: 9383–9387. 10.1073/pnas.1633576100

78. Ashworth L, Aguilar R, Galetto L, Aizen MA. Why do pollination generalist and specialist plant species show similar reproductive susceptibility to habitat fragmentation? J Ecol. 2004; 92: 717–719. 10.1111/j.0022-0477.2004.00910.x

79. Vázquez DP, Aizen MA. Asymmetric specialization: A pervasive feature of plant-pollinator interactions. Ecology. 2004; 85: 1251–1257. 10.1890/03-3112

80. Godínez-Alvarez H, Jordano P. An empirical approach to analysing the demographic consequences of seed dispersal by frugivores. In Dennis A, Schupp EW, Green R, Wescott D editors. Seed Dispersal: Theory and its Application in a Changing World. CAB International; 2007. pp. 391–406. 10.1079/9781845931650.0391

81. Memmott J, Waser NM, Price MV. Tolerance of pollination networks to species extinctions. Proc R Soc Biol Sci. 2004; 271: 2605–2611. 10.1098/rspb.2004.2909

82. de Assis BJ, Guimarães PR, Peres CA, Carvalho G, Cazetta E. Local extinctions of obligate frugivores and patch size reduction disrupt the structure of seed dispersal networks. Ecography, 2018; 41: 1899–1909. 10.1111/ecog.03592

83. McConkey KR, Drake DR. Low redundancy in seed dispersal within an island frugivore community. AoB Plants. 2015; 7: 1–13. 10.1093/aobpla/plv088

84. Reich PB, Tilman D, Isbell F, Mueller K, Hobbie SE, Flynn DFB. Impacts of biodiversity loss escalate through time as redundancy fades. Science 2012; 336: 589–592. 10.1126/science.1217909

85. Cardinale, B. J., Duffy, J. E., Gonzalez, A., Hooper, D. U., Perrings, C., Venail, et al. Biodiversity loss and its impact on humanity. Nature, 2012; 486: 59–67. 10.1038/nature11148

86. Kurek P, Dobrowolska D, Wiatrowska B, Seget B, Piechnik L. Low rate of pre-dispersal acorn predation by eurasian Jays *Garrulus glandarius* during non-mast years. Acta Ornithol. 2023; 57: 211–215. 10.3161/00016454AO2022.57.2.009

87. Ruán I. Dispersión y diversidad genética de *Stenocereus queretaroensis* (Cactaceae) en Jalisco.PhD Thesis. Universidad de Guadalajara. 2016.

88. García-Ruiz, Ruán-Tejeda I, Zuloaga-Aguilar MS, Íñiguez-Dávalos LI. Characterization of endozoochorous dispersal of pitayo *Stenocereus queretaroensis*, in Autlán, Jalisco, Mexico. Ethol Ecol Evol. 2018; 30: 447–460. doi:10.1080/03949370.2017.1423114

89. Cestari C. Bernardi CJ. Predation of the Buffy-fronted seedeater *Sporophila frontalis* (Aves: Emberizidae) on *Merostachys neesii* (Poaceae: Babusoideae) seeds during a masting event in the Atlantic forest. Biota Neotrop. 2011; 11: 407–411. https://10.1590/S1676-06032011000300033

